# Enhancing Challenging Target Screening via Multimodal Protein-Ligand Contrastive Learning

**DOI:** 10.1101/2024.08.22.609123

**Authors:** Zhen Wang, Zhanfeng Wang, Maohua Yang, Long Pang, Fangyuan Nie, Siyuan Liu, Zhifeng Gao, Guojiang Zhao, Xiaohong Ji, Dandan Huang, Zhengdan Zhu, Dongdong Li, Yannan Yuan, Hang Zheng, Linfeng Zhang, Guolin Ke, Dongdong Wang, Feng Yu

## Abstract

Recent advancements in genomics and proteomics have identified numerous clinically significant protein targets, with notably 85% categorized as undruggable. These targets present widespread challenges due to their complex structures and dynamics, rendering conventional drug design strategies not always effective. In this study, we introduce Uni-Clip, a contrastive learning framework that incorporates multi-modal features of proteins (structure and residue) and ligands (conformation and graph). Optimized with a specifically designed CF-InfoNCE loss, Uni-Clip enhances the modeling of protein-ligand interactions for both undruggable and druggable proteins. Uni-Clip demonstrates superior performance in benchmark evaluations on widely acknowledged datasets, LIT-PCBA and DUD-E, achieving a 147% and 218% improvements in enrichment factors at 1% compared to baselines. Furthermore, Uni-Clip proves to be a practical tool for various drug discovery applications. In virtual screening for the challenging protein target GPX4 with flat surface, it identified non-covalent inhibitors with an IC_**50**_ of 4.17 μM, in contrast to the predominantly covalent inhibitors currently known. Through target fishing for benzbromarone, Uni-Clip identified the intrinsically disordered protein c-Myc as a potential target, highlighting benzbromarone’s potential for repurposing in cancer therapy. Explainable analyses effectively identified binding sites consistent with molecular dynamics and experimental results, even for challenging undruggable targets.

## 1 Introduction

With advancements in genomics and proteomics, numerous clinically significant protein targets have been identified in various diseases. These targets can be categorized as druggable (15%) or undruggable (85%) [1]. Undruggable protein targets typically possess complex structures, dynamics, and functions, making them challenging to target using conventional drug design strategies. These challenging targets can be broadly categorized into several groups: proteins with extensive interface areas or relatively flat binding surfaces, such as GPX4, Bcl-2, and MDM2; intrinsically disordered proteins, like c-Myc and Tau; proteins exhibiting high conformational flexibility, including TRIP13 and TG2; and proteins lacking well-defined druggable pockets on their surfaces, as observed in the small GTPase RAS family proteins (e.g., KRAS and HRAS) [2–4]. The high diversity and dynamic characteristics of undruggable targets present substantial challenges in identifying and ranking protein-ligand interactions (PLIs) for these targets.

Due to the strong correlation between protein function and its three-dimensional structure, structure-based methods demonstrate high efficiency and accuracy in modeling PLIs for proteins with experimentally determined structures, particularly druggable proteins [5–7]. However, despite advancements in structure determination technologies such as high-throughput X-ray crystallography and cryo-electron microscopy, high-confidence structures for undruggable proteins remains limited [8, 9]. While artificial intelligence technologies such as AlphaFold can predict protein structures, their confidence levels are relatively low for complex undruggable proteins [10, 11]. In contrast, sequence data for both druggable and undruggable proteins is readily accessible due to high-throughput sequencing technology. Moreover, sequence-based models can avoid biases from structural dynamics such as wiggling and jiggling. Consequently, sequence-based PLI modeling methods have experienced significant development in recent years [12–16]. Rationally combining multiple modalities can effectively integrate the advantages of structure-based and sequence-based methods, providing a flexible framework to improve the screening of complex targets. By doing so, it seeks to more effectively identify new potential targets within the expanding protein space [17–19]. However, developing an efficient approach to integrate multimodal protein information for modeling of PLIs remains a challenge.

Contrastive learning, recognized as a self-supervised learning paradigm, has achieved notable success in the domains of computer vision and natural language processing [20, 21]. Inspired by the success of models like SimCLR [22] and CLIP [23], recent studies have increasingly applied contrastive learning to model PLIs [7, 16, 24]. Unlike regression or classification methods that directly analyze protein-ligand complexes to predict PLIs, contrastive learning encodes proteins and ligands separately and uses a contrastive loss function to enhance the alignment of positive pairs and the differentiation of negative pairs. Through contrastive learning, this approach enhances the efficiency of utilizing PLI affinity data and effectively ranks corresponding candidates for given proteins or ligands. Furthermore, by judiciously selecting positive sample pairs and adjusting the temperature parameter, contrastive learning demonstrates potential robustness against label noise [25].

In this study, we introduce a contrastive protein-ligand pretraining model, termed Uni-Clip. This model is a multimodal contrastive learning framework developed to model protein-ligand interactions (PLIs) that are applicable to both druggable and undruggable protein targets. Uni-Clip concurrently processes protein structure and residue feature, as well as ligand conformation and graph feature, via the integration of diverse pretrained encoders, thereby encompassing the data features handled by other baselines, as shown in Fig. 1c. Subsequently, we have developed the collision-free information noise-contrastive estimation (CF-InfoNCE) loss function, enabling Uni-Clip to more adeptly align the representations of protein-ligand pairs within a unified feature space, substantially enhancing the prediction of their interactions. Furthermore, we constructed the largest Multimodal Protein Ligand Binding (MMBind) dataset to our knowledge, specifically designed to enhance the performance of the Uni-Clip model. We validated the efficacy of Uni-Clip through a practical undruggable protein case target-ing Glutathione Peroxidase 4 (GPX4), identifying DP021 as an effective non-covalent inhibitor with confirmed binding affinity. Uni-Clip also demonstrated its capability in target fishing using benzbromarone, identifying c-Myc as a potential target and showcasing its anti-cancer potential. Our explainable analyses of Uni-Clip effectively identified binding sites consistent with molecular dynamics and experimental results, even for challenging targets with low druggability or intrinsically disordered proteins like c-Myc. We have also shown that Uni-Clip substantially enhances performance compared to previous methods across two benchmark datasets in the field of virtual screening, while also maintaining optimal speed. In summary, this research highlights Uni-Clip as a promising tool not only for achieving strong performance in standard benchmarks, but also for accelerating various drug discovery scenarios.

**Fig. 1:**
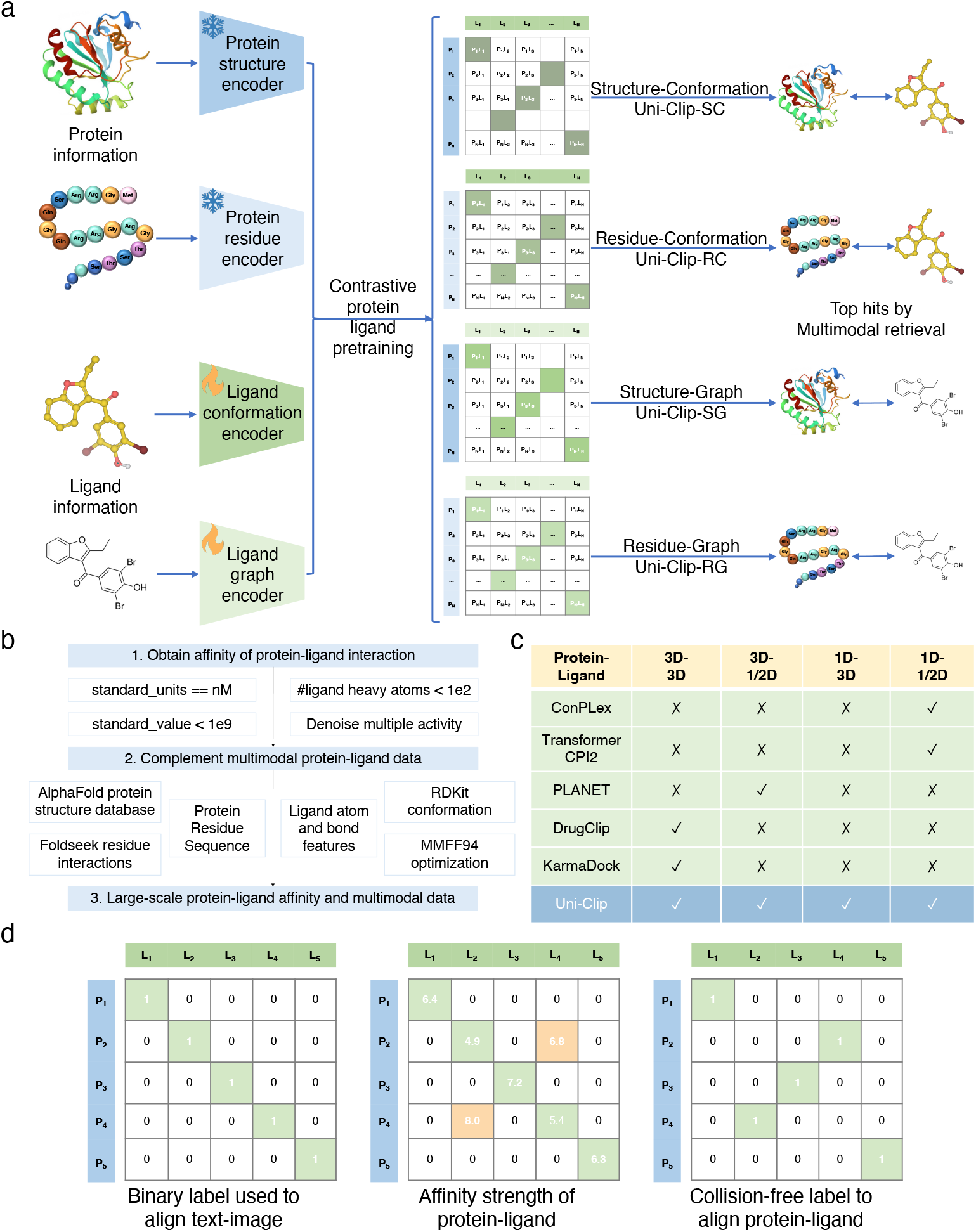
Uni-Clip predicts protein-ligand interactions using multimodal protein-ligand data. **a** The computational pipeline of Uni-Clip. **b** The data curation pipeline of large-scale multimodal protein-ligand data and affinity label. **c** Comparative analysis of input modalities in baselines and Uni-Clip for PLIs. **d** Our proposed collision-free label matrix aligning protein-ligand vs. vanilla binary label used to align text-image.

## 2 Results

### 2.1 AI-driven comprehensive drug discovery platform exploring both undruggable and druggable targets

Previous virtual screening methods in drug discovery, while effective for druggable targets, are limited by their reliance on structural data and struggle with undruggable targets. Methods using protein sequences, though theoretically able to represent traditionally undruggable targets, would benefit from added structural information. It is a critical need for a platform that handles both druggable and undruggable targets, which also boosts the chance to find new targets from expanding protein space for target fishing. To address these issues, we incorporated several enhancements and rectified several critical issues to facilitate both virtual screening and target fishing.

We developed Uni-Clip, comprising approximately 100 million parameters, trained on a dataset of over 850,000 PLI data points. Uni-Clip represents a large-scale model with extensive protein-ligand coverage in the field of PLIs. The effective training of such a model was facilitated by three key components: model design, data curation and training methodology (detail ablation studies of these components are in Supplementary Notes).

The overall framework of Uni-Clip is illustrated in Fig. 1a. There are four distinct encoders: frozen SaProt [26], frozen ESM2 [27], pretrained Uni-Mol [28] and pretrained Uni-Mol-Graph (detailed in Methods 4.1), which respectively encode protein *structure*, protein *residue*, ligand *conformation* and ligand *graph* multimodalities data, which are abbreviated as S, R, C and G. Subsequently, we apply contrastive learning to align the protein structure and residue encoders with the ligand conformation and graph encoders. This alignment leads to four sub-modules: Uni-Clip-SC, Uni-Clip-RC, Uni-Clip-SG and Uni-Clip-RG. By jointly optimizing these four sub-modules, we achieve the final Uni-Clip model, which is adept at handling multimodal features across protein and ligand domains.

We constructed the MMBind dataset from publicly available resources, including the ChEMBL bioactivity database [29] and the AlphaFold protein structure database [30] as depicted in Fig. 1b. The ChEMBL database provides binding affinities of pairs of protein residue sequences and ligand SMILES. We then aligned multimodal data to each protein-ligand pair, including the protein structure data from the AlphaFold protein structure database and ligand conformation and graph data via RDKit. After data quality control, we obtained over 850,000 protein-ligand affinity data points (detailed in Methods 4.2).

Considering that 15% of labels on the off-diagonal dominate over corresponding labels on the diagonal on average during training on the MMBind dataset, we designed a training methodology for protein-ligand contrastive learning with CF-InfoNCE loss, an extension of InfoNCE loss.In contrast, InfoNCE hypothesizes that labels on the diagonal dominate over others in the corresponding row and column, as illustrated on the left side of Fig. 1d. Among the over 850k protein-ligand pairs in the MMBind dataset, there are over 529k ligands but only 5.2k proteins. Consequently a protein can form strong interactions with multiple ligands and vice versa when aligning protein-ligand, as depict in the middle of Fig. 1d. It is important to emphasize that there are two scenarios of collisions in a batch, one is that there may be identical proteins or ligands, and the other is that there may be multiple different ligands (or proteins) binding to the same protein (or ligand). The observed collisions (colored in golden in Fig. 1d) inspired us to develop CF-InfoNCE loss, enabling Uni-Clip to better align a batch of protein-ligand data. To enhance robustness and efficacy from contrastive learning, we only keep the maximal one as shown in the right of Fig. 1d, ensuring that there is one prominent label and significantly smaller remaining values in each row or column. The collision-free label matrix results in row-wise and column-wise normalizations, which separately guide protein-ligand and ligand-protein alignments (detailed in Methods 4.3).

### 2.2 Uni-Clip shows state-of-the-art performance on virtual screening benchmarks

To evaluate Uni-Clip’s performance in virtual screening, we conducted a comparative analysis against several baseline methods: Vina [31], Glide-SP [32], TransformerCPI2 (TCPI2) [15], PLANET [5], and DrugCLIP [7]. Vina and Glide-SP are among the most commonly used molecular docking tools for virtual screening. TCPI2, PLANET, and DrugCLIP are deep learning models designed to utilize different modal representations: 1D sequence-based, 2D graph-based, and 3D structure features, respectively. The efficacy of these tools was benchmarked against two widely acknowledged datasets: DUD-E [33] and LIT-PCBA [34]. To assess screening performance, the Area Under the Receiver Operating Characteristic Curve (AUROC) and enrichment factors (EF_0.5%_, EF_1%_, and EF_5%_) were applied (detailed in Methods Section 4.4). The enrichment factors were calculated as the ratio of true actives in the selected subset versus the entire dataset at sampling rates of 0.5%, 1%, and 5%.

To assess the diversity in retrieval results across different sub-modules of Uni-Clip, we selected the LIT-PCBA dataset for analyzing variances in ligands retrieved by Uni-Clip-SC, Uni-Clip-RC, Uni-Clip-SG and Uni-Clip-RG. We identified the top 1% of ligands by rank from each sub-module for each protein target and computed diversity between the positive ligands. The results revealed that the diversity among the positive ligands retrieved by different retrieval methods exceeded 45% on average (Fig. 2a). This finding suggests that the different retrieval strategies implemented by Uni-Clip sub-modules are highly complementary, effectively enhancing the discovery of a wide range of candidate molecules for the target protein. Despite variations in network architecture and input data types among the encoders in Uni-Clip, we did not impose explicit constraints during the optimization process to manage the divergence between Uni-Clip-SC, Uni-Clip-RC, Uni-Clip-SG and Uni-Clip-RG.

**Fig. 2:**
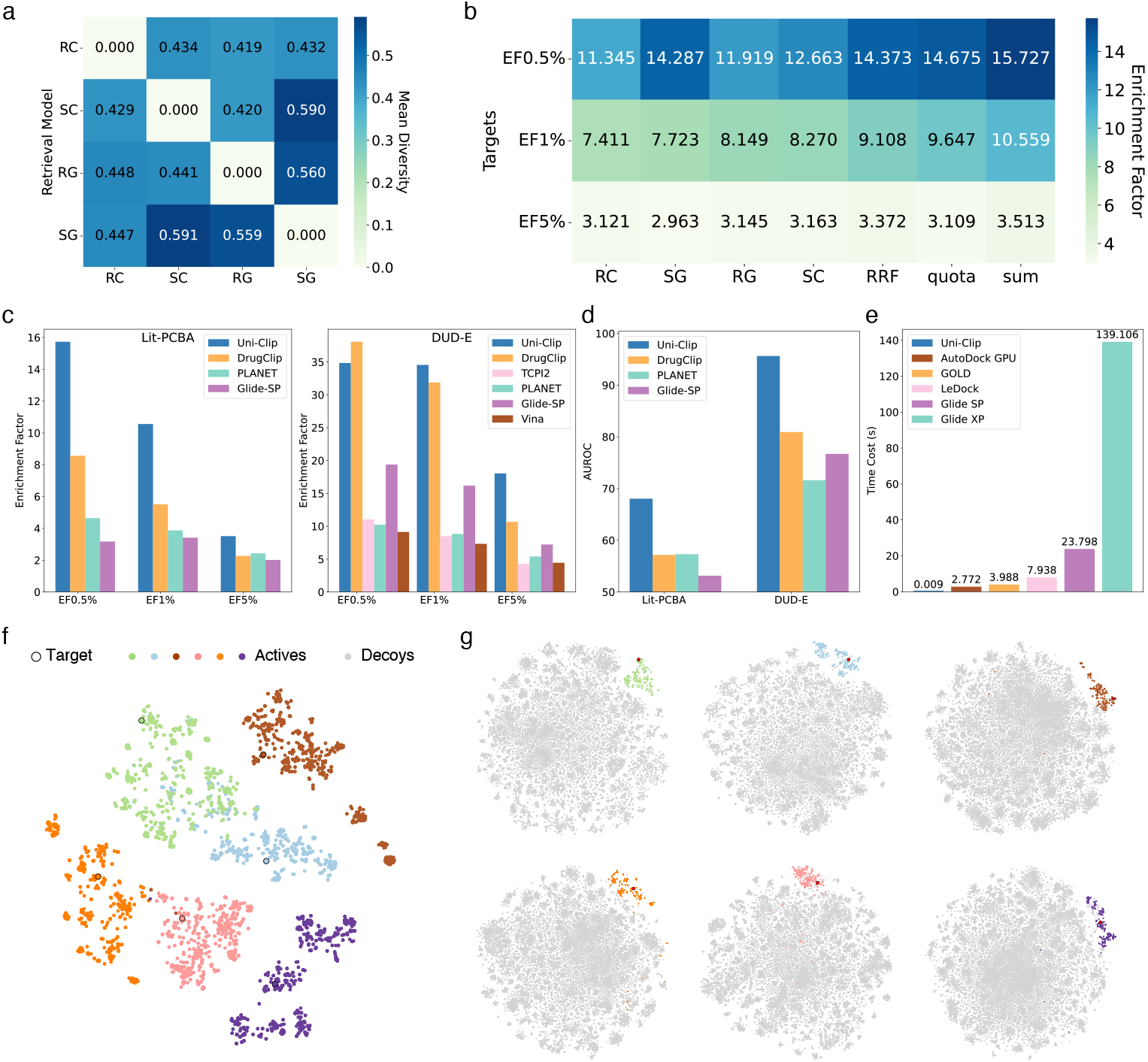
Evaluation of the protein-ligand interactions prediction. **a** Diversity among different multimoldal retrival branches of Uni-Clip. **b** Improvements of Uni-Clip over different multimoldal retrival branches. **c-d** Enrichment factor and AUROC results of Uni-Clip and baseline models on the LIT-PCBA and DUD-E dataset. **e** Average time costs of the various models. **f-g** TSNE visualization of embeddings of DUD-E protein, actives (multicolored) and inactives (gray), produced by Uni-Clip.

Given the variety of retrieval results produced by Uni-Clip sub-modules, we assessed three ensemble strategies to improve the efficacy of Uni-Clip’s virtual screening process: Uni-Clip-sum, Uni-Clip-quota and Uni-Clip-RRF (detailed in the Methods 4.1). As shown in Fig. 2b, on the LIT-PCBA dataset, three ensemble methods outperformed single Uni-Clip sub-module at EF_0.5%_, EF_1%_, and EF_5%_. Particularly, Uni-Clip-sum demonstrated superiority, with an average improvement of 26%, 34%, and 13% compared to the single retrieval method at EF_0.5%_, EF_1%_, and EF_5%_. More detailed results per target are depict in Supplementary Figure A1. In light of the promising performance manifested by Uni-Clip-sum, we employed its score as a representative for Uni-Clip in comparative analyses with other baselines.

As indicated in Fig. 2c, Uni-Clip shows superior performance across most evaluation metrics on the LIT-PCBA and DUD-E datasets. On the LIT-PCBA dataset, the Uni-Clip model outperforms the average performance of all baseline models by 240%, 147%, and 70% at EF_0.5%_, EF_1%_, and EF_5%_, respectively. On the DUD-E dataset, Uni-Clip surpasses the average performance of all baseline models by 162%, 218%, and 216% at EF_0.5%_, EF_1%_, and EF_5%_. Despite not achieving the peak EF_0.5%_ about DUD-E, the result of Uni-Clip remains competitive. More notably, Uni-Clip achieves the highest values for AUROC as shown in Fig. 2d. These comparative improvements highlight the consistency of Uni-Clip’s predictive capabilities across different selection fractions of top-scoring molecules and its potential to substantially increase the effectiveness of virtual screening. Furthermore, to validate the efficiency of Uni-Clip in virtual screening, we illustrate the average time cost of Uni-Clip compared to various docking methods, including Glide-XP [35], Glide-SP [32], LeDock [36], GOLD [37], and AutoDock GPU [38], as shown in Fig. 2e. A lower bar indicates a faster method. In terms of screening power, Uni-Clip demonstrates a considerable advantage over established methods, notably achieving at least a 300-fold acceleration over AutoDock GPU, an already GPU-accelerated docking tool. This further underscores the potential application of Uni-Clip as an efficient tool for ultra-high-throughput ligand-based virtual screening.

For a more intuitive evaluation of the representations produced by Uni-Clip, we utilized t-SNE [39] to visualize the embeddings of Uni-Clip-RC for proteins and ligands from the DUD-E dataset. As shown in Fig. 2f, within the representation space, each target protein is clearly distinguishable, and the corresponding positive ligands display a noticeable clustering pattern. Furthermore, in Fig. 2g, we observed that the representations of positive ligands tend to cluster closely with the target proteins in the embedding space, while the representations of negative ligands are more dispersed and located at a greater distance from the target proteins. This suggests that through contrastive learning, Uni-Clip effectively creates clusters in the representation space, aligning with the protein’s positive and negative samples, thereby enhancing the screening and ranking power for a given protein.

### 2.3 Discovery non-covalent inhibitor targeting Glutathione Peroxidase 4

Glutathione peroxidase 4 (GPX4) is a critically important and challenging drug target. As a crucial antioxidant enzyme within the selenoprotein family [40], it plays a vital role in maintaining cellular redox homeostasis by reducing lipid hydroperoxides and protecting cellular components from oxidative damage [41–44]. Its overexpression in various cancers has been linked to tumor progression and therapy resistance, making GPX4 inhibitors a promising cancer treatment strategy through the induction of ferroptosis and reduction of tumor growth [45, 46]. Currently, most existing GPX4 inhibitors are covalent compounds that utilize electrophilic warheads, such as chloroacetamides [47–51], which may lead to potential issues of selectivity and toxicity. Consequently, there is an urgent demand for potent, drug-like non-covalent small molecule inhibitors of GPX4. However, X-ray crystallography has revealed that the surface of GPX4 is relatively flat and lacks druggable pockets (Fig. 3a and Supplementary Table A3). The highest druggability score observed from the crystal structure of GPX4 is 0.196 on a scale from 0 to 1, where higher values indicate better druggability. This extremely low score presents a significant challenge for traditional structure-based drug discovery (SBDD).

**Fig. 3:**
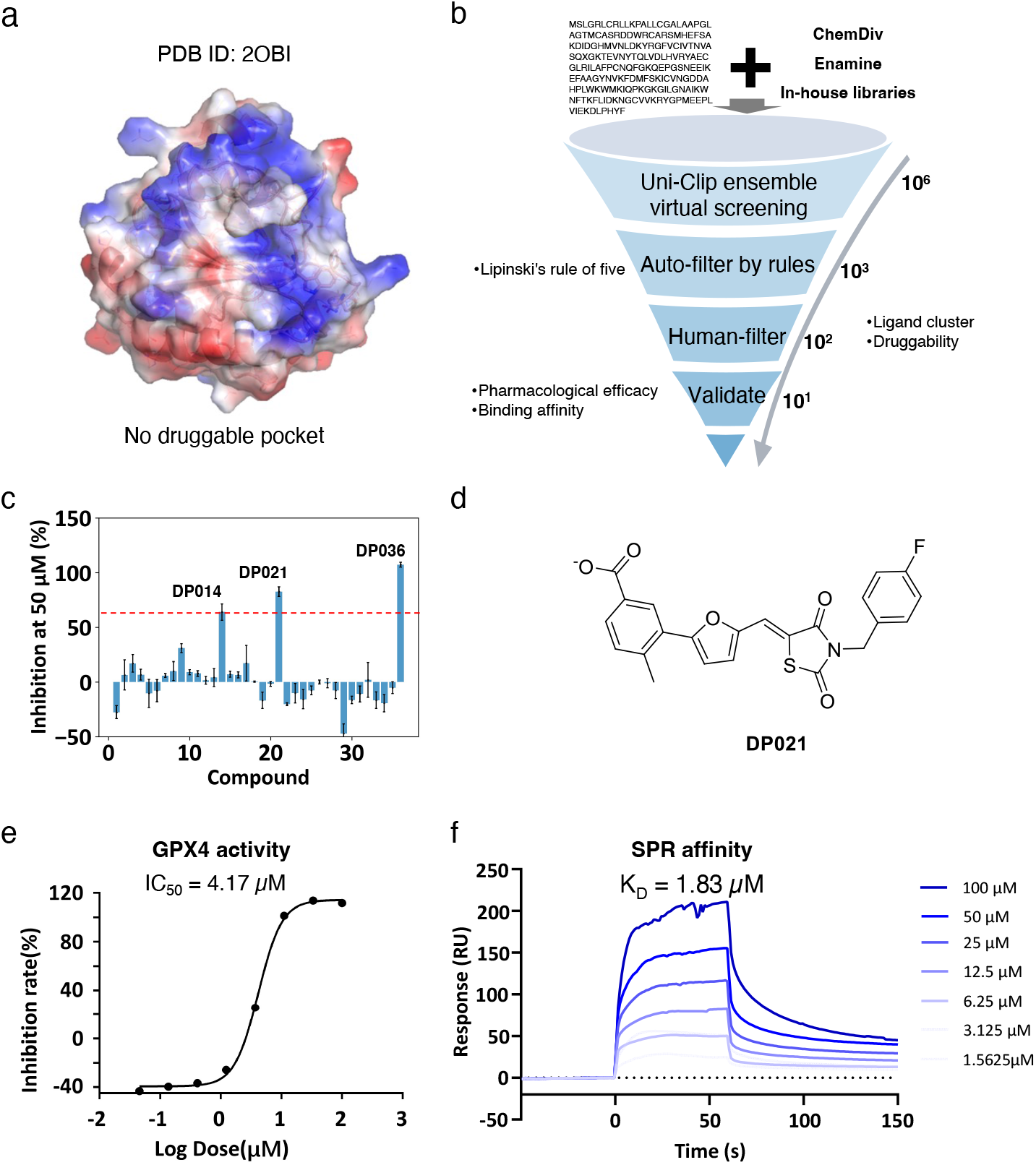
Hit discovering of GPX4. **a** Crystal structure of GPX4 (PDB ID: 2OBI) is depicted, with surface representation. Notably, no druggable pockets are observed in this structure. **b** Schematic representation of the Uni-Clip virtual screening workflow, illustrating the process used to identify potential GPX4 inhibitors. **c** Biological activity inhibition of the 36 candidates at the concentration of 50 *μ*M. **d** Chemical structure of compound DP021, presented in a 2D format. **e** The half-maximal inhibitory concentration (IC_50_) for DP021 against GPX4 enzyme activity is determined to be 4.17 *μ*M. **f** The binding affinity constant (K_*D*_) of DP021 to GPX4, as measured by SPR, is found to be 1.83 *μ*M.

Hence, we employed a multimodal method, Uni-Clip, for the virtual screening of GPX4. Given the protein’s flat surface and the lack of identifiable druggable pockets, it was not clear which of the Uni-Clip sub-modules (Uni-Clip-SC, Uni-Clip-RC, Uni-Clip-SG, Uni-Clip-RG, or Uni-Clip-sum) would be most appropriate. To address this challenge, we devised an ensemble virtual screening protocol that harnesses the five retrieval sub-modules of Uni-Clip, termed “Uni-Clip ensemble virtual screening” (Fig. 3b). We implemented these five approaches to screen and evaluate compounds from two extensive commercial compound libraries, ChemDiv and Enamine, each containing millions of compounds, as well as our in-house compound library comprising approximately 1,900 compounds. Utilizing the five retrieval approaches of Uni-Clip, we selected the top 900 compounds from the commercial libraries and the top 100 from our in-house library, totaling (900 + 100) × 5 = 5, 000 compounds for subsequent analysis. Compounds violating Lipinski’s rule of five or having a druglikeness score [52] below -6 were excluded. The remaining compounds were clustered based on their extended-connectivity fingerprints (ECFP), resulting in 127 distinct chemotype clusters. Representative compounds were selected to ensure structure diversity, and 36 candidates were ultimately chosen for further experimental evaluation.

These compounds underwent an initial assessment of their biological activity through enzyme inhibition assays. Two distinct concentrations, 10 *μ*M and 50 *μ*M, were employed for this purpose. The inhibition rate at 50 *μ*M is depicted in Fig. 3c, while a comprehensive presentation of the results can be found in Supplementary Tables A4. Certain compounds function as inhibitors, exerting a positive effect, while some others seemingly operate as agonists, resulting in a negative impact. Remarkably, the enzyme inhibition rate for three of the compounds exceeded 60% (Fig. 3c). The two compounds exhibiting the highest inhibition rates, DP021 and DP036, were further subjected to multi-concentration fitting analysis. The half-maximal inhibitory concentration (IC_50_) values for DP021 and DP036 were determined to be 4.17 *μ*M (Fig. 3d and 3e) and 26.25 *μ*M (data not shown), respectively. Consequently, DP021 was chosen for more in-depth analysis and experimental testing. Subsequent Surface Plasmon Resonance (SPR) experiments were performed to verify the direct binding interaction between DP021 and the GPX4 protein. The binding affinity constant (K_*D*_) was found to be 1.83 *μ*M (Fig. 3f). To the best of our knowledge, this represents the most potent non-covalent GPX4 inhibitor reported in the literature thus far.

To determine the precise binding site of DP021 on GPX4, we employed alanine scanning mutagenesis within the Uni-Clip framework (refer to Methods) to evaluate the individual contribution of each residue (Δscore) to the binding affinity of DP021 for GPX4 (Fig. 4a). A higher score denotes a greater contribution to the interaction. For alanine scanning analysis, we used the Uni-Clip-RC sub-module, as DP021 was initially identified through this approach. The top 10 residues, identified as potential interactors with DP021, are listed in Fig. 4a. To identify potential binding pockets on the planar surface of GPX4, we performed mixed-solvent molecular dynamics (MixMD) simulations [53] using various small molecules as solvents. A 20 ns production run yielded 1,000 conformations for analysis. Fpocket [54] was subsequently utilized to identify 11,599 pockets. The pockets that contain one or more of the top 10 residues identified by Uni-Clip and have the highest druggability score are listed in Fig. 4b and Supplementary Table A5. Residues 138F, 152R, 52N and 129I were found within the same pocket (Pocket 0), which had the highest druggability score, suggesting this pocket may be the most probable binding site for DP021.

**Fig. 4:**
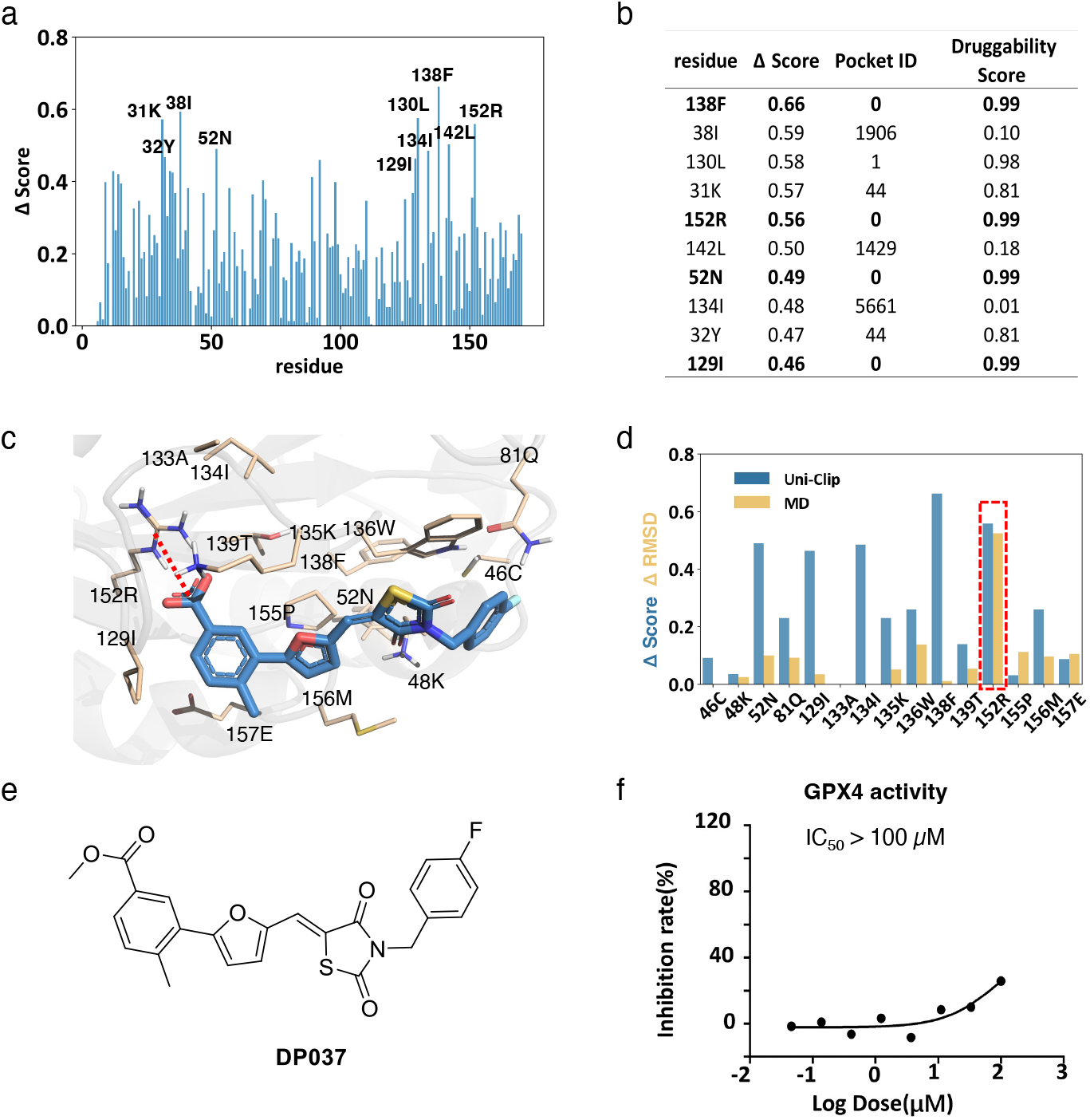
The binding site of DP021 on GPX4. **a** Alanine scanning mutagenesis results from Uni-Clip, illustrating the individual contributions of each residue to the binding affinity of DP021 for GPX4. **b** Pockets of the top 10 residues, as derived from the MixMD. **c** The optimal docking pose of DP021 obtained via induced fit docking. The residues in Pocket 0 are visualized as lines, while DP021 is represented as sticks (PyMol). **d** The specific contributions of each residue in Pocket 0 to the binding of DP021 were evaluated using Uni-Clip alanine scanning (Δ score) and molecular dynamics simulation (Δ RMSD). **e** The chemical structure of the modified compound DP037 is presented in a 2D format. **f** The half-maximal inhibitory concentration (IC_50_) of DP037 on GPX4 enzyme activity.

To validate this hypothesis, we docked DP021 into this pocket using the induced fit docking (IFD) method and selected the docking pose with the highest docking score for further analysis, as shown in Fig. 4c. We mutated each pocket residues of DP021 to alanine and performed molecular dynamics simulations to confirm the key residues in this pocket. The RMSD of the ”wild-type pose” indicates considerable stability (Supplementary Figure A2). For the ”mutated pose,” the RMSD differences (ΔRMSD) from the wild-type pose are used to assess the importance of each residue (Fig. 4d). Notably, the ”mutated pose” of residue R152 is the most unstable (Supplementary Figure A2), with ΔRMSD exceeding 0.5 nm (Fig. 4d), underscoring the critical role of 152R in the interaction between DP021 and GPX4. The docking pose also indicates that the primary interaction arises from a salt bridge formed by residue 152R with the carboxyl functional group of DP021. Therefore, we modified the carboxyl group of DP021 by capping it with a methyl group, creating a novel molecule, DP037 (Fig. 4e). Using Uni-Clip to evaluate DP037, the score decreases by 61% from DP021 to DP037. Subsequent synthesis and IC_50_ testing of DP037 against GPX4 via enzyme activity assays showed that the inhibitory effects of DP037 had considerably decreased with IC_50_ greater than 100uM (Fig. 4f). This result suggests that residue R152 and the formed salt bridge are crucial for DP021 binding, highlighting Pocket 0 as the potential binding site for DP021. This finding also indicates that Uni-Clip can accurately identify the key residues for ligand binding.

In summary, our application of the ”Uni-Clip ensemble virtual screening” successfully led to the identification of DP021 as a GPX4 inhibitor. Additionally, Uni-Clip facilitated the precise identification of the binding pocket and key residues for DP021. Our research demonstrates how Uni-Clip can aid in discovering new inhibitors for challenging targets like GPX4.

### 2.4 Comprehensive target fishing of benzbromarone: exploring undruggable and druggable targets

Target fishing is an emerging approach in drug discovery that aims to identify potential protein targets for a query molecule. This strategy facilitates the clarification of the mechanism of action and biological activities of compounds whose targets remain unknown [55]. In this context, we selected benzbromarone (Fig. 5a), a well-established uricosuric agent and a non-purine selective inhibitor of xanthine oxidase [56], to demonstrate the target fishing capability of Uni-Clip. Benzbromarone is mainly prescribed for hyperuricemia and gout treatment [57], targeting urate transporter 1 (URAT1) to increase renal excretion of uric acid. Despite being approved in several countries for gout management in patients with inadequate response to standard therapy or contraindications to other urate-lowering drugs, benzbromarone’s potential in other therapeutic areas remains largely unexplored.

**Fig. 5:**
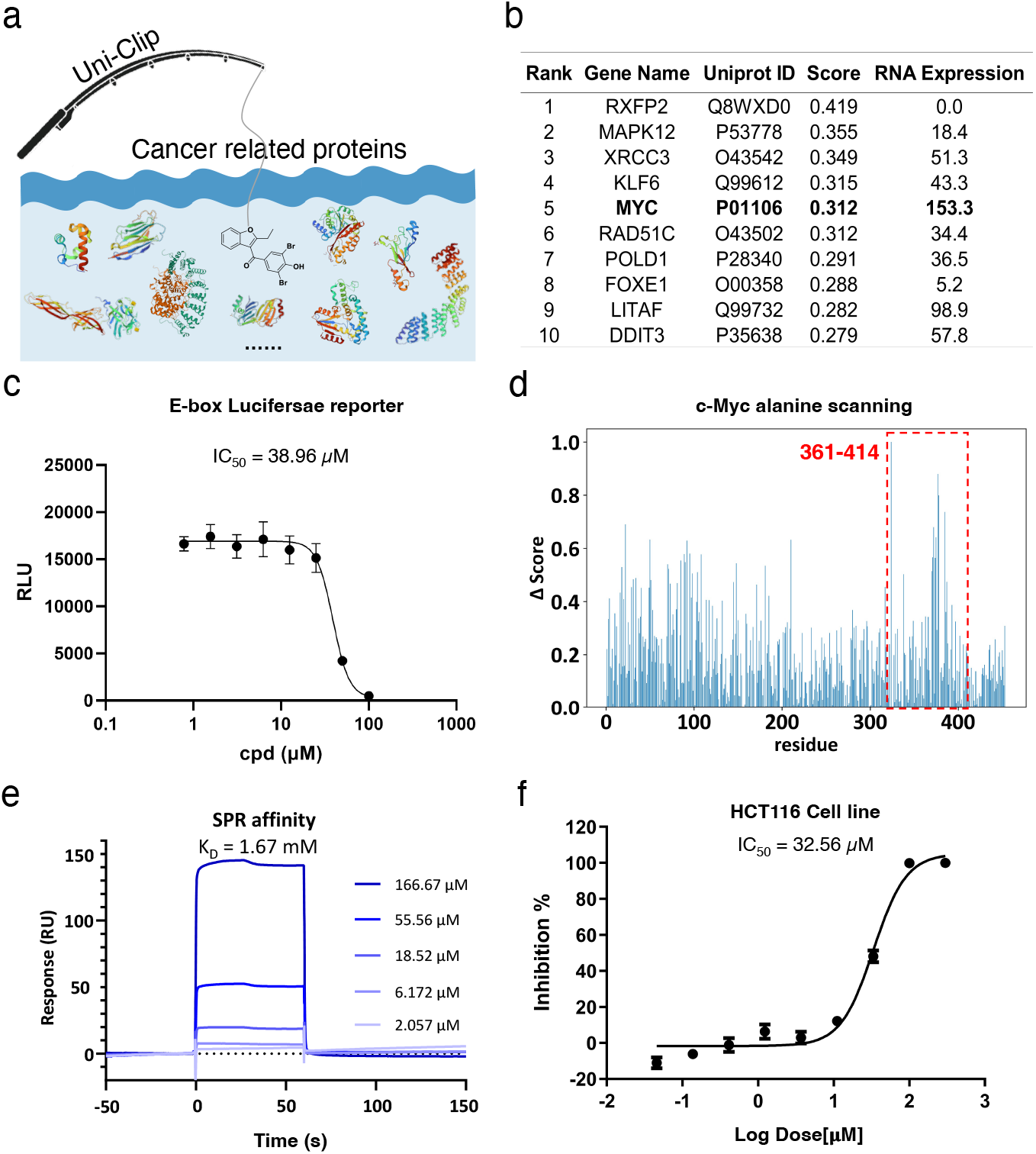
Target Fishing Analysis of benzbromarone. **a** Schematic representation of the Target Fishing strategy utilizing the Uni-Clip. **b** Ranking of the top 10 genes selected by the Uni-Clip-RG approach. RNA expression levels (in nTPM units) were sourced from the Human Protein Atlas database. **c** The c-Myc E-box reporter assay of benzbromarone shows an IC_50_ value of 38.96 *μ*M. **d** Alanine scanning results from the Uni-Clip. The red rectangular dot highlights the binding site of benzbromarone located within the C-terminal domain (residues 361-414) of c-Myc. **e** Surface Plasmon Resonance (SPR) analysis confirming the interaction between benzbromarone and the c-Myc protein, yielding a dissociation constant (K_*D*_) of 1.67 mM. **f** Cellular activity of benzbromarone in HCT116 cells, with an IC_50_ value of 32.56 *μ*M.

Our investigation was prompted by a recent study that utilized machine learning with graph attention networks (GATs) to predict benzbromarone as a potential therapeutic agent for lung adenocarcinoma (LUAD) [58]. Building on this insight, we also used the five retrieval approaches of Uni-Clip (Uni-Clip-SC, Uni-Clip-RC, Uni-Clip-SG, Uni-Clip-RG and Uni-Clip-sum) to evaluate the interaction of benzbromarone with 330 human proteins associated with tumor, as catalogued in the UniProt database (Fig. 5a). The resulting scores were ranked according to the predicted interaction probability. From this analysis, we shortlisted the top 10 proteins from each Uni-Clip method as candidate targets for benzbromarone. To further refine our target selection, we retrieved RNA expression data for lung cancer cell lines from the Human Protein Atlas database [59–62]. Guided by the RNA expression profiles, we identified c-Myc as the primary research target for benzbromarone, which was highlighted by the Uni-Clip-RG method (Fig. 5b). Additional findings of other four retrieval approaches are detailed in Supplementary Table A9.

c-Myc, an intrinsically disordered protein (IDP) and a transcription factor, plays a pivotal role in regulating gene expression and orchestrating a myriad of biological processes [63, 64]. The aberrant expression of c-Myc protein is observed in approximately 70% of tumors, resulting from gene amplification, chromosomal translocation, upregulated mRNA, and abnormal protein stability [63, 65–67]. To investigate the potential of benzbromarone as a c-Myc inhibitor, we employed a systematic approach: first, we verified the benzbromarone’s ability to inhibit c-Myc function; second, we confirmed the benzbromarone’s binding region to c-Myc; and finally, we evaluated the benzbromarone’s effect on related disease cell phenotypes of c-Myc.

To ascertain whether benzbromarone could inhibit c-Myc function, we initially employed a luciferase reporter assay utilizing an E-box element (Fig. 5c). Our findings indicate that benzbromarone exerts an inhibitory effect on c-Myc function, with a half-maximal inhibitory concentration (IC_50_) of 38.96 *μ*M.

To elucidate the direct binding of benzbromarone to c-Myc and identify the binding region, we utilized the alanine scanning approach of Uni-Clip (Fig. 5d). This analysis localized the binding sites to the C-terminal domain (residues 361-414) of c-Myc. We then synthesized a corresponding c-Myc peptide spanning this region and conducted a binding assay using SPR to evaluate the direct interaction with benzbromarone. The SPR data demonstrates a dose-dependent binding of benzbromarone to the c-Myc protein on the biosensor chip, with a dissociation constant (K_*D*_) of 1.67 mM, confirming the direct binding of benzbromarone to the c-Myc C-terminal domain (residues 361-414).

Further investigations were conducted to evaluate the impact of benzbromarone on the cancer cells, specifically examining its effects on HCT116 cells. Benzbromarone demonstrates inhibitory effects on the proliferation of HCT116 cells, with an IC_50_ value of 32.56 *μ*M (Fig. 5f). These findings collectively suggest that benzbromarone may exert anti-cancer effects through its direct interaction with c-Myc, thereby inhibiting cancer cell growth.

This study underscores the potential of benzbromarone as a therapeutic agent targeting the c-Myc protein, offering a promising avenue for cancer treatment through drug repurposing. The ability of benzbromarone to directly bind to and inhibit the transcriptional activity of c-Myc highlights its potential utility in combating cancers characterized by dysregulated c-Myc expression.

## 3 Discussion

Structure based virtual screening methods in drug discovery have traditionally been effective for druggable targets but face limitations when dealing with undruggable targets. This challenge stems from the methods’ reliance on structural data, which is often scarce or unreliable for complex, undruggable proteins. To address this issue, we developed Uni-Clip, a multimodal contrastive learning model aimed at enhancing the modeling of PLIs for both druggable and undruggable proteins. By combining for pretrained encoders for both proteins and ligands, Uni-Clip effectively integrates protein structure and residue data with ligand conformation and graph data through the CF-InfoNCE loss, resulting in a comprehensive representation of protein-ligand interactions. Consequently, Uni-Clip consists of four submodules: Uni-Clip-SC, Uni-Clip-RC, Uni-Clip-SG, and Uni-Clip-RG. The multimodal design of Uni-Clip, in conjunction with the multimodal PLI dataset we constructed, MMBind, enhances its versatile applicability in diverse drug discovery scenarios, such as virtual screening and target fishing.

Uni-Clip effectively addresses challenging cases by better handling both druggable and undruggable proteins, offering a more comprehensive solution for virtual screening. Compared to traditional single-modal approaches, Uni-Clip offers significant advantages by leveraging multimodal data to improve predictive accuracy and computational efficiency on widely acknowledged datasets LIT-PCBA and DUD-E. In the actual drug discovery process, Uni-Clip has effectively identified DP021 as a prospective inhibitor targeting the challenging protein GPX4, with its inhibitory efficacy corroborated by experimental assays. Furthermore, within a comprehensive database of druggable and undruggable proteins, Uni-Clip identified the binding interaction between benzbromarone, an established uricosuric agent, and an intrinsically disordered oncogenic protein, c-Myc, thereby highlighting benzbromarone’s potential as an anticancer therapeutic. Moreover, utilizing the alanine scanning method, Uni-Clip demonstrates potential in identifying binding sites within real-world applications.

While Uni-Clip has shown promising results, it still has potential for further refinement. First, as a data-driven neural network model, Uni-Clip’s generalizability can be greatly improved by expanding the PLI dataset to cover a broader range of protein and ligand distributions. Additionally, collecting more high-quality experimental data and leveraging data derived from first-principles calculations are crucial steps for enhancing its performance. Second, although Uni-Clip has been effective in utilizing multimodal data of PLIs, incorporating more detailed information, such as protein binding pockets and ligand fragments, could enhance both the model’s effectiveness and flexibility. In addition, Uni-Clip offers multimodal encoders for proteins and ligands. In practical applications, when addressing diverse protein-ligand pairs, incorporating a Mixture of Experts module would facilitate the dynamic selection and weighting of outputs from different encoders based on the input features of proteins and ligands. This adaptive mechanism would enable Uni-Clip to flexibly adjust to various protein-ligand combinations for more complex undruggable proteins. Despite these challenges, Uni-Clip represents a substantial advancement in PLI modeling, offering enhanced accuracy and efficiency, with wide applications in virtual screening and target fishing, whether for druggable or undruggable proteins. Uni-Clip could be a valuable tool for medicinal chemists, with great potential to accelerate the drug discovery process.

## 4 Methods

This section offers a comprehensive summary of each component in the proposed method and provides details of the experiments. We elucidate the key characteristics of our workflow and configurations, encompassing model architecture design, data curation, evaluation metrics, and procedures for experiments conducted both in silico and in the wet lab.

### 4.1 Encoders for Uni-Clip

We use individual encoder for each of the modality for both ligands and proteins in Uni-Clip. For ligands, which are essentially small molecules, we use Uni-Mol as the structure encoder and develop a new method Uni-Mol-Graph as the molecular 2D-topology encoder. For proteins in 1D sequence and 3D structure, we chose ESM2 and SaProt, respectively. Here we briefly introduce these modules. Uni-Mol is a robust 3D representation model for molecules. It has been pretrained on 209 million molecule conformations using unsupervised learning, achieving state-of-the-art results in various molecular representation learning tasks [28]. Uni-Mol integrates atomic and coordinate features to capture the three-dimensional structure of ligands by a two-track transformer architecture. To capture key components in molecule 2D-graph such as chemical bonds and avoid fluctuated results given by 3D-encoder for molecules with high flexibility in comformational space, we developed Uni-Mol-Graph, a molecule encoder leveraging only features within the molecular graph(shortest path between nodes, for instance). Uni-Mol-Graph shares the same model backbone as Uni-Mol but excludes inter-atomic distance expansions from its attention mechanism and requires only a 2D molecular graph as input. Detailed input features were specified in Supplementary Table A7 and Table A8. Similar to Uni-Mol, Uni-Mol-Graph was pretrained on the Uni-Mol dataset using a masked atom tokens recovery strategy. After the pretraining phase, we employed Uni-Mol-Graph as the graph encoder for ligands. Both Uni-Mol and Uni-Mol-Graph were finetuned during the PLI training process.

Evolutionary scale modeling 2 (ESM2) is chosen as protein residue encoder. ESM2 is a sequence-based protein language model proposed by the Meta Fundamental AI Research(FAIR) Protein Team. It is pretrained on 250 million protein sequence data in a masked language model manner. Among sequence-only methods, ESM2 stands out as one of the most formidable competitors. The 650M version of ESM2 is used in this research. Since fine-tuning a model at this scale is computationally expensive, we detach ESM2 from the training process and save ESM2 representations for all sequences in advance.

For protein structure, we use the recently proposed SAProt as the encoder. SAProt integrates both structure tokens and residue tokens at residue level in protein language model based on the architecture of ESM2, which enables it to be aware of protein structure information. Pretrained on 40 million protein structures generated by AlphaFold, SaProt shows strong capacity in term of protein representation [68].

Atom-level representations from molecule encoders and residue-level representations from protein encoders are mean-pooled to form molecule- and protein-level representations. Given the multimodal representations, a projection head with sparse activation function *s* [69] is applied to each of them to align the semantic space as:

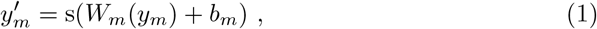

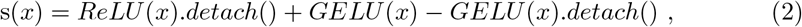

*m* refers to a certain modality. Specifically, the sparse activation function *s* gives a neuron ReLU outputs but GELU [70] gradient to avoid the dead neuron problem. Activated by ReLU, the representations become rather sparse, which greatly facilitates the calculation of cosine similarity between modalities.

To synthetically consider four results given by the combination of different encoders, we applied a summation ensemble strategy, where all candidates are reranked according to the summed score from all results. Since equal weights is assigned to each part of the loss function corresponding to the combination above during training, we keep their weights equal when conduct the ensemble:

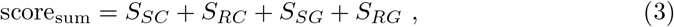

score_sum_ is the score of certain pair in sum ensemble, *S*_*SC*_, *S*_*RC*_, *S*_*SG*_, *S*_*RG*_ represents the score given by Uni-Clip-SC, Uni-Clip-RC, Uni-Clip-SG, Uni-Clip-RG, respectively. The ensemble result is termed as Uni-Clip-sum. We also tried quota fusion strategy and reciprocal rank fusion (RRF) strategy, termed as Uni-Clip-quota and Uni-Clip-RRF (Supplementary Notes ), but the results shows that simple summation strategy has the best performance on benchmarks. Consequently, we use only this strategy through out remaining experiments.

### 4.2 MMBind Dataset

The construction of MMBind dataset includes the collection of protein-ligand affinity values and preparation of multi-modal ligand and protein data. We achieve the first goal by cleaning and refining ChEMBL33, a large database of bioactive assays collected from published experimental data, providing ligand canonical SMILES, protein UniProt IDs and their assay results. Initially, the original ChEMBL dataset was first filtered by assay type of ’B’ (Binding), target type of ’SINGLE PROTEIN’ and molecule type of ’Small molecule’. All samples with assay type of Ki, Kd, IC50 and EC50, which occupies over 99.6% of all samples, were included for further processing while others were dropped. Moreover, data with abnormal concentration values were further removed, such as negative and extremely large ones. To make full use of the dataset, a threshold of 562 *μM* molar concentration was set to define positive samples. Additionally, we filtered out molecules with more than 100 heavy atoms. To achieve a zero-shot test, all targets in the DUD-E and LIT-PCBA datasets were excluded. Given the UniProt IDs of the targets, we accessed protein sequences using API provided by UniProt. The UniProt IDs was also used to access the AlphaFold database to get 3D fold structure. Although some of the proteins had experimental crystalline structure, we chose to include only AlphaFold structures in consistency with SAProt. For small molecules, 3D conformations needed by Uni-Mol were randomly generated and optimized using the MMFF94 force field by RDKit. Molecule topology was also obtained in this procedure. After these steps, we constructed a dataset containing 853,932 pairs composed of 533,656 different ligands and 5245 proteins, which, to our knowledge, is one of the largest datasets in field of PLIs as shown in Supplementary Table A6. In this research, the MMBind dataset is randomly split at a ratio of 10:1 for training and validation.

### 4.3 Collision-free InfoNCE loss

The collision-free InfoNCE loss is designed to solve the multiple-positive-samples issue in context of PLI. It incorporates an dynamic labeling strategy into the conventional InfoNCE loss, which writes:

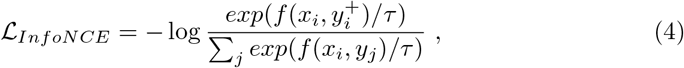

in which *x*_*i*_ is the representation of sample *i*, 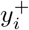 is the corresponding positive sample’s representation, and *τ* is a temperature hyperparameter. *f* is a scoring function measuring the similarity between representations. Since the available experimental data contains majorly positive samples of binding proteins and ligands, construction of negative pairs is of necessity. Similar to CLIP, we adopt a dynamic in-batch sampling strategy. Given a batch of *N* pairs 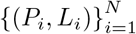 , the combination of 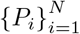 and 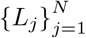 results in *N* ^2^ pairs (*P*_*i*_, *L*_*j*_).

Usually, only pairs with *i* ≠ *j* are treated as positives samples while others are negative ones. However, as shown in Fig. 1d, it is possible that the dynamic constructed off-diagonal (*i* ≠ *j*) pairs turn to be positive samples. We refer this issue as “collision of positive samples” and try to solve it with the following dynamic labeling strategy.

Specifically, each pairs (*P*_*i*_, *L*_*j*_) is first assigned with a label *l*_*i*,*j*_ related to its assay value:

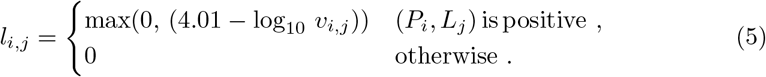

where *v*_*ij*_ is the assay value of pair (*P*_*i*_, *L*_*j*_), For each row in the resulting label matrix, the pair with largest label, i.e., the pair with strongest binding affinity, is selected as the positive sample, while other pairs are regarded as negative samples. After that, the label matrix is normalized row-wise. In this way, the label is made sure to be one-hot and the dominant pair is preserved, avoiding the collision problem. The collision-free InfoNCE loss ℒ_𝒞ℱ_ can be written as:

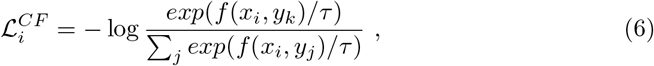

where *k* = arg max_*j*_ *l*_*ij*_. Considering the protein and ligand parts, the overall loss function of a batch data for a retrieval branch *B* ∈ (*SC, RC, SG, RG*) is:

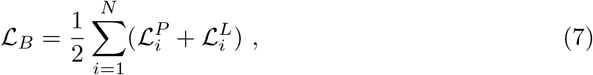

where ℒ_ℬ_ is the InfoNCE loss calculated with collision-free label. In Uni-Clip, we align 4 different representations: the SaProt and ESM2representations for proteins with the Uni-Mol and Uni-Mol-Graph representations for ligands. Their combination, i.e. 4 contrastive pairs *L*_*SC*_, *L*_*RC*_, *L*_*SG*_ and *L*_*RG*_, forms the final loss as:

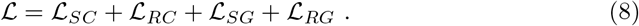

### 4.4 Evaluation metrics and datasets

Evaluation metrics includes AUROC and enrichment factor (EF) on 0.5%, 1% and 5%. While AUROC is widely used in classification tasks, enrichment factor is a metric for virtual screening, measuring how good a screening method performs compare to random guess:

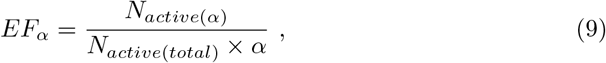

where *α* is a proportion, *N*_*active*(*α*)_ is the number of true active compounds in the top *α* candidates sorted by model. *N*_*active*(*total*)_ is the total number of active compounds among all candidates.

Diversity between candidates sets is obtained from different retrieval branches. This metric is calculated from the fingerprints’ similarity between the molecules they contain. The diversity between set *A* against set *B* is:

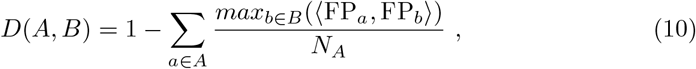

*D* is diversity value, FP is the Morgan fingerprint of a molecule, *N*_*A*_ is the number of elements of set *A*.

DUD-E and LIT-PCBA have been chosen to evaluate Uni-Clip’s screening power. DUD-E is one of the most popular virtual screening benchmark datasets [71–73]. It contains 102 proteins. Each protein is associated with a known ligand defining the binding pocket and relevant binders as well as non-binders in a ratio of 1:50.

LIT-PCBA is another virtual screening benchmark dataset proposed to address biased data problem in previous benchmarks [34]. It is built upon dose-response Pub-Chem bio-assays, consisting of 15 targets with 7,844 experimentally active and 407,381 inactive compounds.

### 4.5 Identifing important functional groups and residues

As shown in results, Uni-Clip can not only handle scoring and ranking tasks, but also help determine the key interactions in the protein-ligand pairs. This is achieved through two analysis tools: substitution effect analysis and alanine scanning. Below, we briefly introduce these methods.

In substitution effect analysis, the protein is fixed and the effect of substituting functional groups in its active compounds is examined. Specifically, potential key functional groups in these compounds are substituted with less chemically active ones and the modified molecules’ affinity scores with the protein are subsquently evaluated. The model is expected to give lower scores and recognize the activity cliffs in these molecules.

Alanine scanning is a commonly accepted method to identify important residues in a protein sequence. For proteins, we anchor an active compound and mutate each residue into alanine, which usually does not interact with small molecules due to its lack of a functional group but has similar contributions to protein structure as other amino acids. We check whether the scores predicted by Uni-Clip for the original protein and mutated protein change substantially as Δ*score* = |*score*_*mutation*_ − *score*_*wt*_|.

### 4.6 Virtual screening of GPX4

We leveraged five retrieval approaches of Uni-Clip (Uni-Clip-SC, Uni-Clip-RC, Uni-Clip-SG, Uni-Clip-RG and Uni-Clip-sum) to assess and score potential compounds from three comprehensive chemical libraries. These included ChemDiv, with an inventory of 1,566,730 compounds, Enamine, housing 2,056,498 compounds, and our proprietary library, which contains approximately 1,900 compounds. From the commercial libraries, we selected the top 900 compounds from each method, complemented by the top 100 from our in-house collection, amassing a total of 5,000 compounds for further analysis.

Our objective was achieved through a meticulous two-tier filtering process. Initially, compounds contravening Lipinski’s Rule of Five (a criterion for assessing the drug-likeness of compounds) were eliminated. Simultaneously, we applied DataWarrior software [52] to exclude molecules with a computed drug-likeness score below -6.

Subsequently, the remaining compounds underwent automated clustering based on their ECFP. This molecular descriptor encapsulates the structure attributes of the compounds, leading to the identification of approximately 127 unique clusters, each representing a distinct chemotype.

From these clusters, we meticulously selected representative compounds based on their intra-cluster ranking and their structure diversity relative to other highly ranked compounds. This strategy was designed to ensure a broad spectrum of structure diversity within the final candidate pool. Ultimately, 36 candidates were procured for subsequent experimental validation.

### 4.7 GPX4 protein expression and purification

The GPX4 (residues 28-197) gene was cloned into the pCDNA3.4 vector and coexpressed with SECISBP2 in HEK293 cells, as previously described [74, 75]. The cells were then harvested and resuspended in lysis buffer, containing 50 mM Tris-HCl (pH 7.5), 500 mM NaCl, 10% glycerol, 1 mM dithiothreitol (DTT), and 0.1 mM phenylmethylsulfonyl fluoride (PMSF). The cells were subsequently lysed through sonication, and the cell debris was removed by centrifugation to obtain the supernatant. Next, the supernatant was applied to a Ni-NTA resin column to isolate the protein of interest via affinity chromatography. The column was washed, and the target protein was eluted. To further purify the protein, size-exclusion chromatography was performed using a Superdex 200 column, pre-equilibrated with SEC buffer composed of 50 mM Tris-HCl (pH 7.5), 150 mM NaCl, and 10% glycerol. Finally, the purified protein was aliquoted, flash-frozen in liquid nitrogen, and stored at -80 °C for future use.

### 4.8 GPX4 activity assay

An enzymatic activity inhibition assay was conducted to evaluate the inhibitory potential of selected compounds on GPX4. This assay was performed using the commercially available Cayman GPX4 Inhibitor Screening Assay Kit (catalog number 701880) as per the manufacturer’s instructions. The assay is based on the principle of measuring GPX4 activity indirectly through a coupled reaction with glutathione reductase (GR), where the oxidation of NADPH to NADP+ and the subsequent decrease in absorbance at 340 nm is proportional to GPX4 activity. This colorimetric method allowed us to assess the inhibitory effects of various compounds on GPX4 by monitoring the changes in absorbance. We conducted a preliminary screening of our 36 candidate molecules to evaluate their inhibitory effects, using two different concentrations: 10 *μ*M and 50 *μ*M. Following the identification of the most promising candidates, we determined their half-maximal inhibitory concentration (IC_50_) values. To obtain these values, we performed dose-response experiments by treating the target with increasing concentrations of the selected inhibitors and monitoring the inhibitory response. This comprehensive evaluation of the candidate molecules enabled us to select the most effective inhibitors for further in-depth studies.

### 4.9 GPX4 SPR

The GPX4 SPR biosensing experiments were conducted using a Biacore T200 instrument (GE Healthcare). The running buffer for immobilization was prepared using phosphate-buffered saline (PBS, pH 7.4) supplemented with 5% DMSO, 750 mM NaCl, and 0.05% TWEEN80, which is referred to as PBST. The GPX4 (residues 28-197) protein was immobilized onto a CM5 sensor chip (GE Healthcare), with immobilization levels reaching approximately 3900 response units (RUs). Interaction studies were carried out in PBST at a constant temperature of 25°C. The test molecules were injected for 60 seconds at a flow rate of 50 *μ*L/min, and the dissociation phase was monitored for up to 240 seconds. Data processing and analysis were performed using the Biacore T200 evaluation software (GE Healthcare). The data were fitted to a single-site binding model, enabling the calculation of dissociation constants (K_*D*_ values).

### 4.10 Induced fit docking and molecular dynamics simulations

#### 4.10.1 Mixed-solvent molecular dynamics simulations

Due to the planar surface of GPX4, we employed a mixed-solvent molecular dynamics (MixMD) [53] simulation approach to investigate the potential formation of pockets within the protein. Five small fragments were selected as solvents: acetate ion, ethyl-dimethyl-ammonium ion, 9-methyl-9H-purin-6-amine, N-methylformamide, and 2-methylpyridine. Each solvent was mixed with water at a 1:100 ratio. The AMBER14SB force field [76] was used for the protein, while the GAFF2 [77, 78] force field was applied for the solvent molecules. The TIP3P water model was used for water molecules. A cubic box with a side length of 7.10 nm was constructed, containing a total of 25,394 atoms. Molecular dynamics simulations were performed using the OpenMM [79] software package. The system was first energy minimized, followed by equilibration in the NVT ensemble for 100 ps and then in the NPT ensemble for 100 ps to ensure system stability. Finally, a 20 ns production run was conducted in the NPT ensemble, with conformations saved every 10 ps, resulting in a total of 1,000 frames for subsequent analysis. Fpocket [54] software was used to analyze the pockets in the obtained 1,000 conformations, yielding a total of 11,599 pockets. We identified the pocket with the highest druggability score and contains one of the top 10 residues as the pocket of that residue.

#### 4.10.2 Induced fit docking

We employed the IFD method to investigate the binding interactions between small molecules and the GPX4 protein. Due to the relatively flat surface of GPX4, traditional rigid docking approaches may not accurately capture the conformational changes induced by ligand binding. In contrast, the IFD method allows for flexibility in both the ligand and the protein receptor, thereby providing a more realistic representation of the binding process. The IFD protocol was carried out using the Hermite web platform at https://hermite.dp.tech. For each small molecule, multiple docking poses were generated and evaluated based on their IFD scores, which take into account both the binding affinity and the conformational energy of the protein-ligand complex. The top-scoring pose was selected as the representative binding conformation for further analysis.

#### 4.10.3 Conventional molecular dynamics simulations

To validate the importance of the pocket residues identified for the DP0221 molecule, we performed mutagenesis and conventional molecular dynamics (MD) simulations on the GPX4-DP021 complex obtained from the IFD analysis. Protein mutations were introduced using the protein preparation tool in the Hermite web platform at https://hermite.dp.tech. The pocket residues were individually mutated to alanine (Ala). The force field parameters used for the MD simulations were consistent with those employed in the mixMD approach. The simulations were carried out using the uni-GBSA automated simulation tool [80], with each mutation being simulated for 300 ns. To account for the stochastic nature of MD simulations, three parallel replicas were run for each mutation. For trajectory analysis, the 50-100 ns segment of each trajectory was considered. The root-mean-square deviation (RMSD) values of the small molecule relative to its initial conformation were calculated for all conformations sampled after 50 ns in each of the three replicas. The average RMSD value for each mutation was then computed and subtracted from the corresponding value for the wild-type (WT) complex. This difference in RMSD was used as a stability score for each mutation site, reflecting the impact of the mutation on the binding stability of DP021.

### 4.11 Cell lines

HCT116 cells (ATCC, CCL-247) were cultured in McCoy’s 5A medium (16600-082, Gibco), while HEK293T cells (catalog no. CBP60439, Cobioer) were maintained in Dulbecco’s Modified Eagle Medium (DMEM; catalog no. C0162-811S, CellWorld). Both culture media were supplemented with 10% (v/v) heat-inactivated fetal bovine serum (FBS; catalog no. 10099141C, Gibco) and 1% (v/v) penicillin-streptomycin solution (P/S; catalog no. C0160-611, CellWorld) to support optimal cell growth and viability.

### 4.1.2 c-Myc E-box reporter assay

HEK293T cells were seeded onto a 6-well plate at a density of 1 × 10^6^ cells per well, following harvesting and counting. After a 6-hour incubation period, which allowed the cells to adhere to the plate, a transfection mixture was prepared. This mixture contained 16 *μ*L of FuGENE® HD Transfection Reagent (E2311, Promega) and 4 *μ*g of Ebox-Luciferase plasmid (11544ES03, YEASEN). The components were mixed thoroughly and incubated for 15 minutes before being added to each well. The cells were then cultured in a 37°C incubator with 5% CO_2_ for an additional 18 hours.

Subsequently, transfected cells were seeded onto a 96-well plate at a density of 3 × 10^4^ cells per well. Compounds were added simultaneously to each well at various concentrations (50 *μ*M maximum concentration, double dilution series, 10 concentration gradients), with DMSO serving as a positive control. Three replicates were performed for each well. After a 24-hour incubation period, the cell supernatant was removed, and luminescence was measured using the Luciferase Reporter Gene Assay Kit (11401ES80, YEASEN), following the manufacturer’s instructions.

Specifically, 100 *μ*L of lysis buffer was added to the cells, and samples were incubated for 5 minutes on ice. Next, 20 *μ*L of cell lysate was transferred to a white 96-well plate (410096M, in vitro scientific) and combined with 100 *μ*L of Firefly luciferase test reagent. Luminescence was then measured to assess luciferase activity. Additionally, 20 *μ*L of the cell lysate was transferred to a black 96-well plate (J09602, JingAn biological), where 20 *μ*L of CellTiter-Glo (G9242, Promega) was added to determine cell viability through luminescence measurements. Luminescence values were normalized by the corresponding relative cell viability (CTG value of each well/mean CTG value of each compound).

### 4.13 c-Myc SPR

The c-Myc SPR assay was performed utilizing a c-Myc peptide (residues 361-414) synthesized by Hefei Guotai Biology Science and Technology. The peptide was immobilized onto a CM5 sensor chip, which had been pre-treated with a 1:1 mixture of 0.4 M 1-ethyl-3-(3-dimethylaminopropyl)carbodiimide (EDC) and 0.1 M N-hydroxysuccinimide (NHS). The immobilization level achieved was 6192 response units (RU). Subsequently, the surface of the CM5 chip was blocked with an ethanolamine solution for 10 minutes to eliminate non-specific binding. The compound of interest was prepared at the desired concentration in a buffer containing 5% dimethyl sulfoxide (DMSO) and phosphate-buffered saline with Tween 20 (PBST). The Biacore T200 instrument (GE Healthcare) was employed to determine the association (K(a)) and dissociation (K(d)) constants of the compound, enabling the calculation of its binding affinity. The equilibrium dissociation constant (K_*D*_) was computed using the steady-state model and a 1:1 binding kinetic model.

### 4.14 Proliferation assay

HCT116 cells were seeded in a 96-well plate at a density of 500 cells per well. Concurrently, compounds were added to each well at varying concentrations, ranging from 300 *μ*M as the maximum concentration, with a triple dilution series consisting of 9 gradients. DMSO-treated wells were included as positive controls, and each condition was replicated twice. The cells were then incubated for 6 days at 37°C in a 5% CO_2_ atmosphere.

Following incubation, 50 *μ*L of CellCounting-Lite 2.0 Luminescent Cell Viability Assay reagent (G7573, Promega) was added to each well. The plates were gently shaken for 2 minutes, after which the relative luminescence units (RLU) were measured to determine cell viability. The inhibition rate was calculated using the following formula:

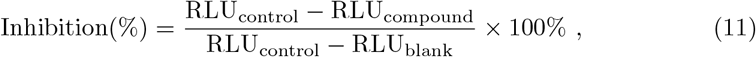

where RLU_compound_ represents the RLU values for compound-treated wells, RLU_blank_ denotes the RLU values for DMSO wells without cells, and RLU_control_ signifies the RLU values for positive control wells.

## 5 Data availability

ChEMBL database is available at https://www.ebi.ac.uk/chembl/. The commercial Enamine Library is available at https://enamine.net/, and ChemDiv Library is available at https://www.chemdiv.com/. All data are available from the corresponding author upon request. The source data can be accessed through figshare at https://figshare.com/s/cb245e12c48a2298d610.

## 6 Code availability

The source code will be available after paper acceptance. Several servers, such as virtual screening, targt fishing and binding site prediction, are available as a non-commercial usage at https://bohrium.dp.tech/apps/uni-clip/job?type=app.

## 7 Acknowledgements

This study was supported by the National Key Research and Development Program of China (2022YFA1004302).

## Appendix A Supplementary Notes, Tables and Figures

Here is the supplementary ablation study of Uni-Clip. To further substantiate the hypothesis that multimodal training can enhance PLI performance, we conduct experiments that training Uni-Clip on single-modal and multimodal PLI data. All experiments are trained on the MMBind dataset and evaluated on the LIT-PCBA dataset. The experiment results of multimodal training are summarized in Supplementary Table A1, multimodal training outmatches any single-modal training on the LIT-PCBA dataset, with an average improvement of 40%, 30%, and 8% at EF_0.5%_, EF_1%_, and EF_5%_.

As Supplementary Table A2 show, CF-InfoNCE surpasses vanilla InfoNCE on the LIT-PCBA dataset, with an improvement of 15% and 14% at EF_0.5%_ and EF_1%_. Before scoring PLIs, non-negative reparameterization like ReLU, can be operated to protein and ligand features, inducing good feature sparsity and robustness. But ReLU output features may suffer from the dead neuron problem during training. So we adopt straight-through Relu to resurrecte the dead neurons while ensuring sparsity (detailed in Methods 4.1). The Supplementary Table A2 reveals the sparse activation function can obtain an improvement of 4% and 13% at EF_1%_, and EF_5%_ on the LIT-PCBA dataset. We also conduct an ablation study to evaluate the pretrained models: pretrained Uni-Mol, pretrained Uni-Mol-Graph. Results are summarized in Supplementary Table A2. We can see that performance improves by adding each of these pretrained models, with an averaged improvement of 16% and 30% at EF_0.5%_ and EF_1%_.

The quota fusion strategy and RRF strategy are another two ensemble strategies, as mentioned in main text. Unlike Uni-Clip-sum, they are based on rank of pairs rather than their scores. In quota fusion, the candidates with highest scores from each combination is took out to form a retrieval set. This operation is repeated until the set achieves given capacity.

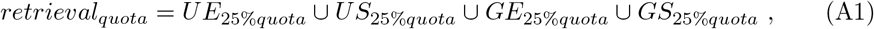

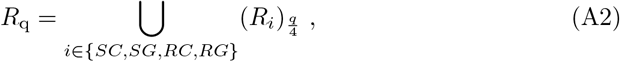

in which *R*_q_ stands for the retrieved set at a given quota q of the candidate set. The ensemble retrieval results are then sorted according to their Uni-Clip-RC scores.

RRF strategy ensembles different ranking results base on a scoring funcion of rank:

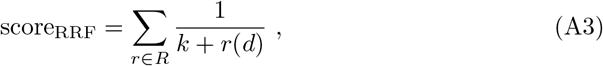

in which *r*(*d*) is the rank of *d* in a ranking result *R, k* is a hyper-parameter and is set to 60 in this reasearch. High score_RRF_ means the candidate is favored in RRF ensembling.

**Table A1:**
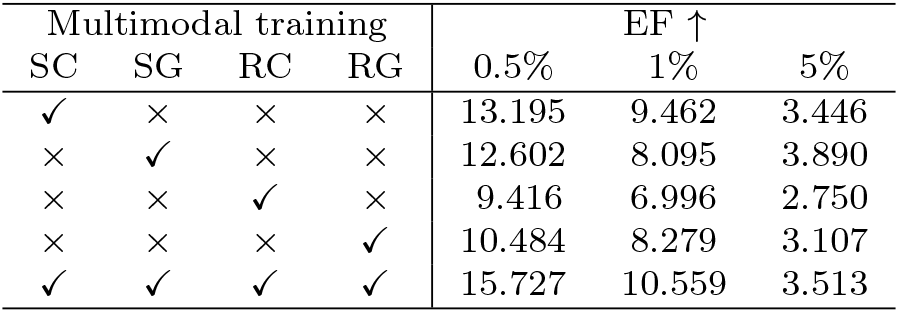
Multimodal training.

**Table A2:**
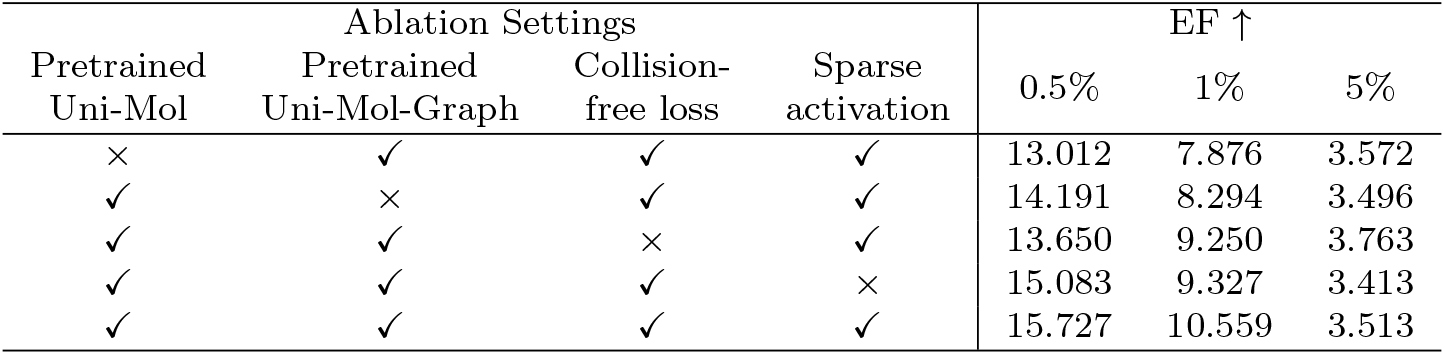
Ablation Settings.

**Table A3:**
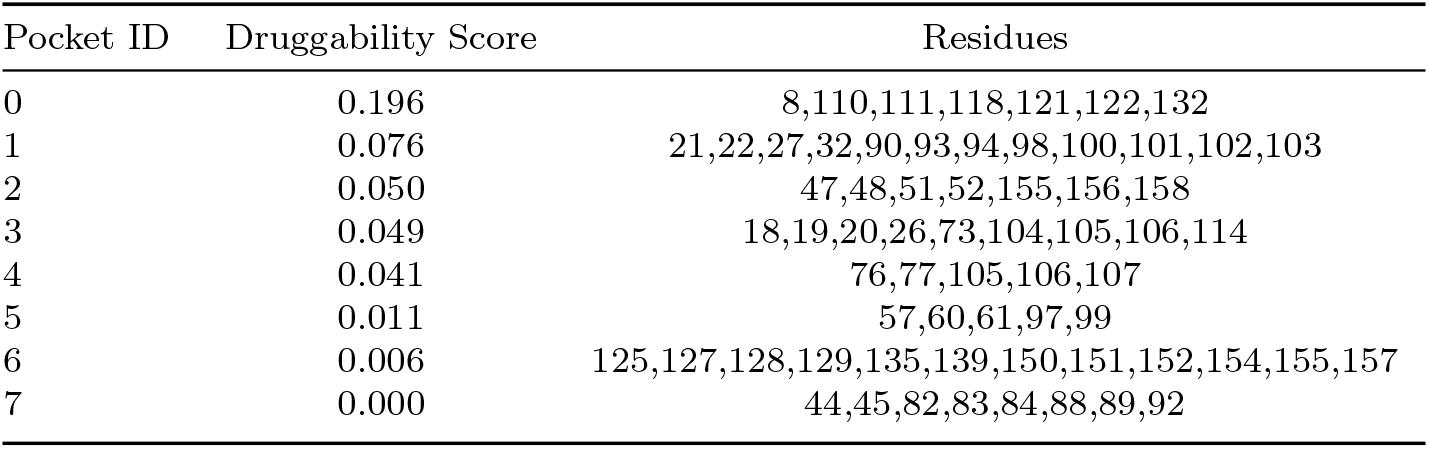
Pockets found in the cystal structure (PDBID: 2OBI).

| Pocket ID | Druggability Score | Residues |
| --- | --- | --- |
| 0 | 0.196 | 8,110,111,118,121,122,132 |
| 1 | 0.076 | 21,22,27,32,90,93,94,98,100,101,102,103 |
| 2 | 0.050 | 47,48,51,52,155,156,158 |
| 3 | 0.049 | 18,19,20,26,73,104,105,106,114 |
| 4 | 0.041 | 76,77,105,106,107 |
| 5 | 0.011 | 57,60,61,97,99 |
| 6 | 0.006 | 125,127,128,129,135,139,150,151,152,154,155,157 |
| 7 | 0.000 | 44,45,82,83,84,88,89,92 |

**Table A4:**
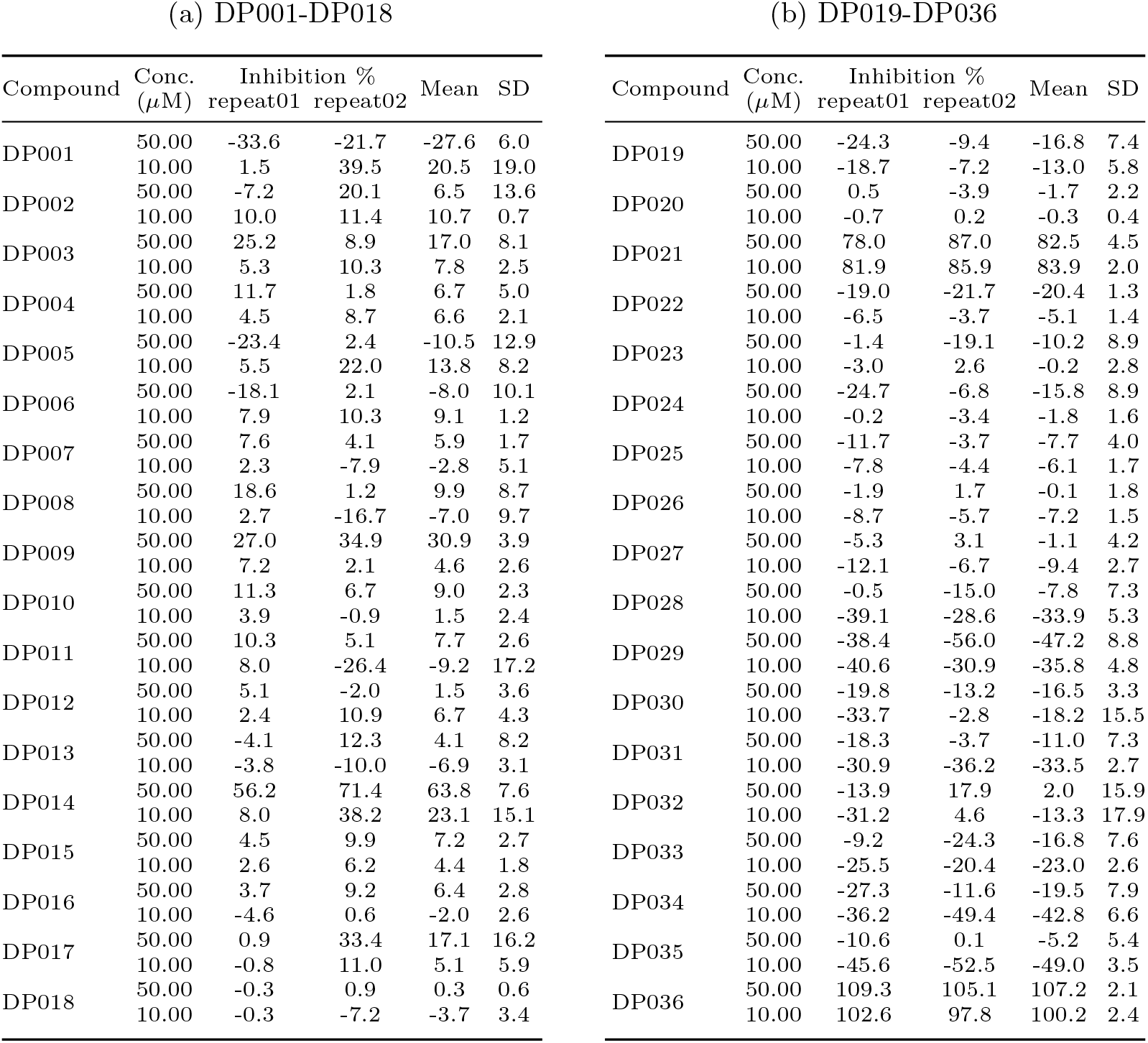
Enzymatic activity inhibition assay results of 36 compounds (DP001-DP036).

**Table A5:** Pockets found in the mixMD structure which contains the top 10 key reidues.

| Pocket ID | Druggability Score | Residues |
| --- | --- | --- |
| 0 | 0.988 | 46,47,48,**52**,81,125,127,128,**129**,135,136,137,**138**,139,150,151,**152**,153,154,155,156 |
| 1 | 0.981 | 46,47,48,81,127,128,129,**130**,135,136,137,139,151,152,153,154,155,156,157,161,164,165 |
| 44 | 0.808 | 21,22,23,27,28,29,**31,32**,52,87,90,91,93,94,98,100,101,102,103 |
| 1429 | 0.175 | 125,**142**,150,151,164,165,168,169 |
| 1906 | 0.100 | **38**,56,59,68,70,162,166 |
| 5661 | 0.005 | 130,131,133,**134**,135 |

**Table A6:** Constuction of training data in several learning methods.

|  | Database | Training Samples |
| --- | --- | --- |
| Uni-Clip | ChEMBL33 | 853932 |
| TCPI2 | ChEMBL27 | 217,732 |
| ConPlex | BindingDB/DUD-E<sup>1</sup> | 12,668/415,204 |
| PLANET | PDBbind 2020/ChEMBL29<sup>2</sup> | 15,616/161,547 |
| DrugCLIP | PDBbind 2019/ChEMBL | 758,107<sup>3</sup> |

<sup>1</sup>ConPlex is trained on BindingDB and fine-tune on DUD-E
<sup>2</sup>Decoys are collected from ChEMBL while binders are from PDBbind
<sup>3</sup>DrugCLIP proposes a data augmentation method, which greatly expands the training set

**Table A7:** Atom features.

| features | size | description |
| --- | --- | --- |
| atom type | 119 | type of atoms including C, N, O, etc, by atomic number |
| chirality | 6 | type of chirality like Tetrahedral chirality |
| degree | 11 | the degree of an atom in molecule |
| formal charge | 11 | integer electronic charge assigned to atom |
| number of H | 9 | number of bonded hydrogen atoms |
| number of radical electrons | 5 | number of radical electrons |
| hybridization | 5 | SP, SP2, SP3, SP3D, SP3D2 |
| aromaticity | 1 | whether an atom is part of an aromatic system |
| in ring | 1 | whether an atom is within a ring structure |

**Table A8:** Bond features.

| features | size | description |
| --- | --- | --- |
| bond type | 4 | SINGLE, DOUBLE, TRIPLE, AROMATIC |
| stereo | 6 | NONE, Z, E, CIS, TRANS, ANY, |
| conjugated | 1 | whether the bond is conjugated, |
| SPD | 512 | the shortest distances between each pair of connected atoms by the Floyd-Warshall algorithm[81] |

**Table A9:** Top ten genes selected by (a) Uni-Clip-SG model, (b) Uni-Clip-SC model, (c) Uni-Clip-sum model and (d) Uni-Clip-RC model.

(a) Uni-Clip-SG
| Rank | Gene Name | Uniprot ID | Score | RNA Expression |
| --- | --- | --- | --- | --- |
| 1 | MSR1 | P21757 | 0.306 | 2.6 |
| 2 | MAPK12 | P53778 | 0.305 | 18.4 |
| 3 | RAD51 | Q06609 | 0.304 | 20.9 |
| 4 | CD274 | Q9NZQ7 | 0.292 | 7.4 |
| 5 | SLC25A11 | Q02978 | 0.288 | 64.1 |
| 6 | HNF1B | P35680 | 0.275 | 2.8 |
| 7 | XRCC3 | O43542 | 0.266 | 51.3 |
| 8 | TBXT | O15178 | 0.265 | 8.2 |
| 9 | TRIP13 | Q15645 | 0.262 | 62.3 |
| 10 | CHCHD7 | Q9BUK0 | 0.261 | 46.5 |

(b) Uni-Clip-SC
| Rank | Gene Name | Uniprot ID | Score | RNA Expression |
| --- | --- | --- | --- | --- |
| 1 | MSR1 | P21757 | 0.380 | 2.6 |
| 2 | CD274 | Q9NZQ7 | 0.363 | 7.4 |
| 3 | RAD51 | Q06609 | 0.325 | 20.9 |
| 4 | MAX | P61244 | 0.319 | 47.1 |
| 5 | XRCC3 | O43542 | 0.315 | 51.3 |
| 6 | MSLN | Q13421 | 0.298 | 28.4 |
| 7 | RAD51C | O43502 | 0.296 | 34.4 |
| 8 | CRTC1 | Q6UUV9 | 0.294 | 9.6 |
| 9 | HNF1B | P35680 | 0.294 | 2.8 |
| 10 | DDIT3 | P35638 | 0.292 | 57.8 |

(c) Uni-Clip-sum
| Rank | Gene Name | Uniprot ID | Score | RNA Expression |
| --- | --- | --- | --- | --- |
| 1 | XRCC3 | O43542 | 0.327 | 51.3 |
| 2 | RXFP2 | Q8WXD0 | 0.323 | 0.0 |
| 3 | MAPK12 | P53778 | 0.312 | 18.4 |
| 4 | RAD51C | O43502 | 0.306 | 34.4 |
| 5 | KLF6 | Q99612 | 0.293 | 43.3 |
| 6 | MSR1 | P21757 | 0.292 | 2.6 |
| 7 | RAD51 | Q06609 | 0.288 | 20.9 |
| 8 | SLC25A11 | Q02978 | 0.287 | 64.1 |
| 9 | CD274 | Q9NZQ7 | 0.286 | 7.4 |
| 10 | DDIT3 | P35638 | 0.278 | 57.8 |

| Rank | Gene Name | Uniprot ID | Score | RNA Expression |
| --- | --- | --- | --- | --- |
| 1 | RXFP2 | Q8WXD0 | 0.456 | 0.0 |
| 2 | XRCC3 | O43542 | 0.376 | 51.3 |
| 3 | RAD51C | O43502 | 0.369 | 34.4 |
| 4 | PARN | O95453 | 0.362 | 22.0 |
| 5 | KLF6 | Q99612 | 0.361 | 43.3 |
| 6 | POLD1 | P28340 | 0.355 | 36.5 |
| 7 | **MYC** | **P01106** | **0.344** | **153.3** |
| 8 | CREB1 | P16220 | 0.343 | 20.3 |
| 9 | MAPK12 | P53778 | 0.324 | 18.4 |
| 10 | DDX6 | P26196 | 0.320 | 44.4 |

**Fig. A1:**
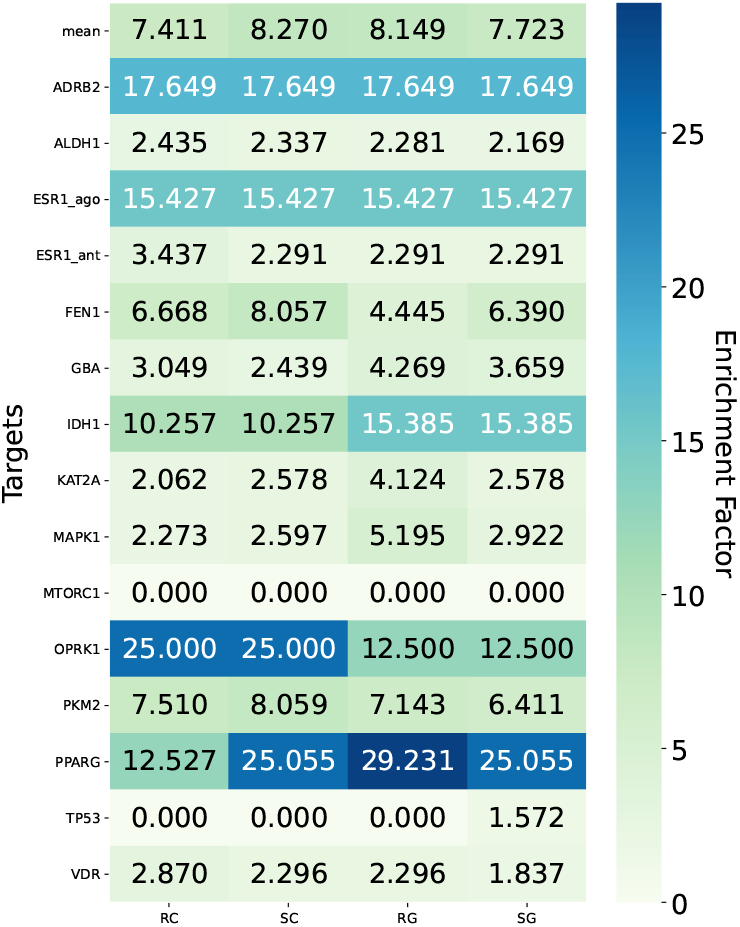
Impact of multimoldal protein ligand data on the EF_1%_ performance of Uni-Clip for different targets on the LIT-PCBA dataset.

**Fig. A2:**
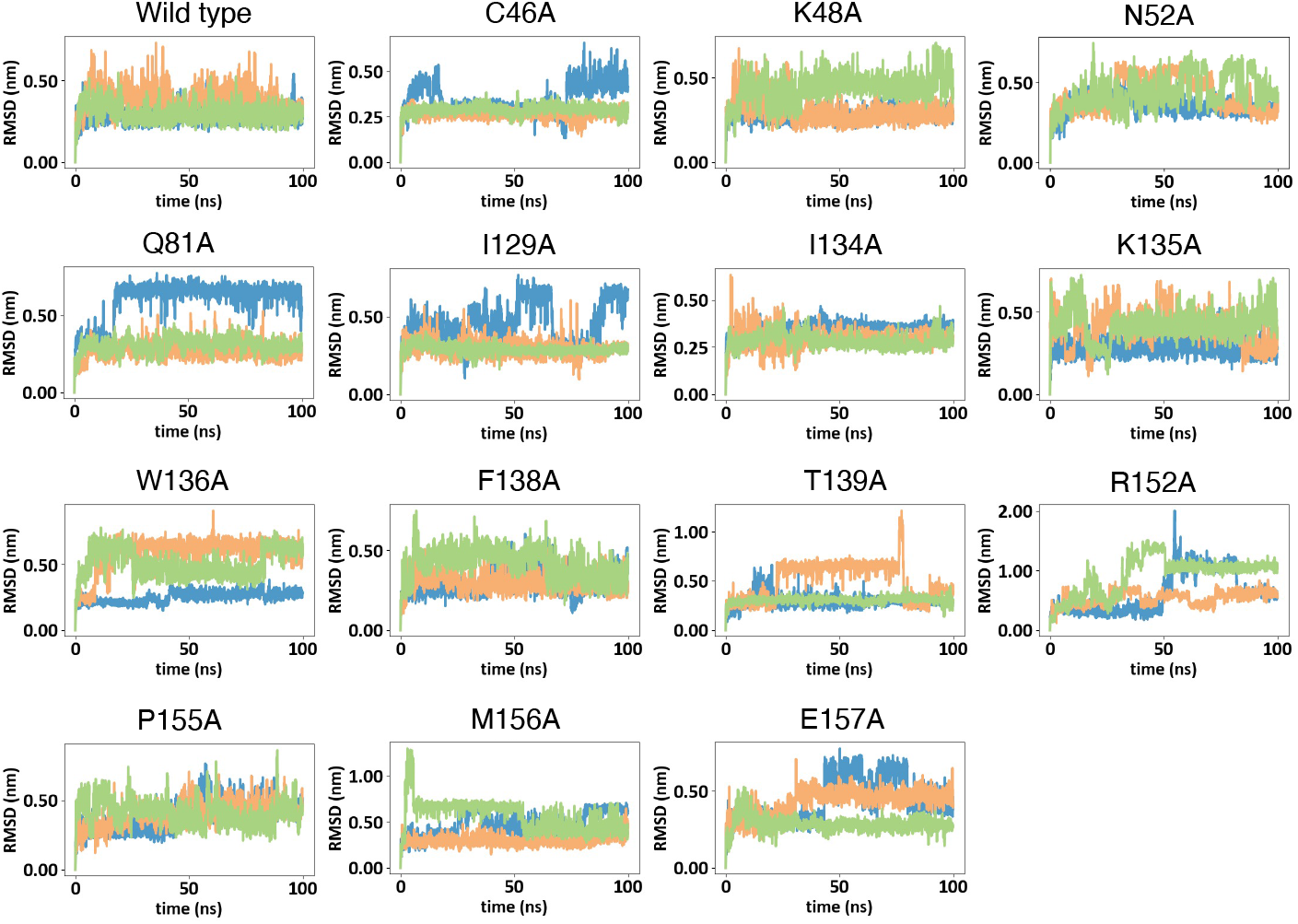
Line plot of RMSD versus time for different mutant protein complexes with DP021, where different colors represent different replicates.

